# Incorporation of ChiA into AcMNPV occlusion bodies and its implications for oral infectivity

**DOI:** 10.64898/2025.12.24.696386

**Authors:** Cinthia Ayelén López, Ricardo Salvador, Gabriela Barros, Antonio Brun, María del Pilar Plastine, Victoria Alfonso, Oscar Taboga, María Gabriela López

**Author notes:** These authors contribute equally to this work (Cinthia Ayelén López, Ricardo Salvador).

## Abstract

Baculoviruses are insect-specific viruses primarily employed in biological pest control and recombinant protein expression. As biopesticides, they offer numerous advantages over chemical pesticides and hold significant potential for application in integrated pest management (IPM). However, their slower action compared to conventional synthetic strategies has limited their widespread adoption. This study aimed to investigate the impact of incorporating a viral protein, AcChiA chitinase, which disrupts the gut peritrophic membrane of susceptible insects, into AcMNPV occlusion bodies *in trans*. This was achieved through the development of an insect cell line that facilitates the passive incorporation of this protein. The presence of ChiA was detected in dissolved occlusion bodies (OBs) preparations from infected transgenic lines, in a method-dependent manner, with the stronger signals observed in chimeric polyhedra. It is hypothesized that ChiA in OBs will increase peritrophic membrane permeability and facilitate primary infection. In bioassays, OBs produced in ChiA-expressing lines were associated with higher larval mortality than OBs produced in Sf9, consistent with a ChiA-mediated effect. Notably, the addition of ChiA proteins from stably transformed insect cell lines improved infection rates, particularly in recombinant baculoviruses, which exhibited a significant reduction in occlusion-derived virion (ODV) occlusion. This underscores the critical role of these proteins in the oral infection process. Consequently, it was possible to incorporate the ChiA protein from stably transformed insect cell lines into wild-type and recombinant baculovirus occlusion bodies, paving the way for potential applications in IPM with AcMNPV.

## Introduction

The use of chemical pesticides has several drawbacks, necessitating the implementation of integrated pest management (IPM) strategies to significantly reduce or eliminate their use. Biological control methods, such as natural enemies and insect-specific pathogens, make a significant contribution to these strategies (Haase et al., 2015). Baculoviruses are the most commonly used viruses for biological control due to their ability to produce occlusion bodies (OBs) and their high specificity, which often affects only one insect species (Cory and Evans, 2007). These viruses are considered safe for vertebrates, as they do not infect mammals. Baculoviruses specifically infect insect larval stages, making them a useful addition to other agents that act at different stages in the pest life cycle for effective crop protection (Abd-Alla et al., 2020). Throughout their life cycle, baculoviruses exhibit two morphologically distinct phenotypes: Occlusion-Derived Virions (ODV) and Budded Virions (BV). The ODVs initiate the primary infections of intestinal larval cells. The reason for this denomination is that they are embedded into a crystalline matrix, shaping the occlusion bodies (OBs) or polyhedra, which act as resistance structures for the virus in the environment and which contaminate the leaves on which the larvae feed (Slack and Arif, 2006). Once ingested, OBs are transported to the midgut, where they are dissolved by the combined action of the alkaline environment and proteases, leading to the release of the ODVs. Once the ODVs infect intestinal epithelial cells, the other virion phenotype, the budded viruses (BVs), spreads the infection towards other cell types. In a very late stage of the infection cycle, polyhedrin (POLH) becomes the protagonist by dramatically increasing its expression level, thereby giving rise to polyhedra in the nucleus. These crystalline structures are highly stable and resist solubilization, except under strong alkaline conditions. Therefore, polyhedra can protect ODVs from most environmental conditions (Rohrmann et al., 2008). For other environmental factors, such as UV, prevention can be achieved through the formulation of the biological insecticide, as suggested by Anggraini et al. (2022).

Strategies to increase the speed of action of insecticides based on recombinant baculoviruses (BVr) typically involve inserting a heterologous gene into the virus genome (Kroemer et al., 2015). During viral replication within host cells, the transgene product (typically a toxin or physiological effector) is expressed alongside the virus proteins (Reid et al., 2023). This gene expression leads to paralysis and disruption of insect homeostasis. BVr also incorporates proteins into OBs as a translational fusion to POLH, which is highly effective among other baculovirus modifications (Kamita et al., 2005). POLH is the primary protein found in OBs. It was determined that an additional copy of wild-type POLH was necessary to properly incorporate fusion proteins into OBs without affecting their morphogenesis by altering the POLH structure (Je et al., 2003). Incorporating toxins into polyhedra is one of the most effective approaches for achieving lethal doses and death times of target insects (Chang et al., 2003). These developments also considered the potential risks associated with releasing genetically modified organisms into the environment (Tiedje et al., 1989).

Chitinases are a diverse group of enzymes that can directly degrade chitin to low-molecular-weight oligomers. They are present in a wide range of organisms, including viruses, bacteria, fungi, yeasts, plants, actinomycetes, arthropods, and humans (Adrangi and Faramarzi, 2013). In 1995, Hawtin et al. identified a functional chitinase gene in the genome of Alphabaculovirus aucalifornicae (ex AcMNPV) (AcChiA). This gene is expressed during the late phase of virus replication in insect cells, where high levels of endo- and exo-chitinase activity were measured. The protein has been detected in both OBs (Hawtin et al., 1995) and in BV (Wang et al., 2010). AcCHIA acts together with a protein called cathepsin, degrading the chitin of the insect exoskeleton and causing liquefaction of the host larva for better dispersal of viral progeny (Hawtin et al., 1997; Ishimwe et al., 2015). According to Rao et al. (2004), the expression of ChiA in bacteria was found to be effective in disrupting the intestinal peritrophic membrane of *Bombyx mori*. This disruption could increase the infectivity of the baculovirus. The objective of this work was to develop a strategy for enhancing the oral infectivity of AcMNPV baculovirus by incorporating viral protein AcChiA into its occlusion bodies, as these proteins have been shown to have disruptive effects on the peritrophic membrane of susceptible larvae. Sf9 lines were developed with the ChiA gene, with and without the addition of the nuclear localization signal KRKK. ChiA was detected in polyhedra samples obtained from infections of both transgenic lines with wild-type AcMNPV and recombinant reporter GFP baculoviruses. The proposed strategy is distinguished by its innovative integration of the ChiA protein into the polyhedra, a process that is not inherent to the POLH structure. This approach constitutes a valuable tool for obtaining wild-type baculovirus inocula with improved capacities. Furthermore, since the enzyme is not fused to polyhedrin, there is no risk of generating recombinant viruses when the occlusion bodies are obtained.

It can also be used for generating double recombinant polyhedra. The potential synergistic effect between ChiA and other proteins, such as toxins or hormones, could enhance the bioinsecticide effect of baculoviruses during primary infection. However, in bioassays, ChiA enhanced the infectivity levels of both wild-type and recombinant baculovirus. Moreover, its addition improved infection in the particular case of recombinant baculoviruses that showed a notable reduction in ODV occlusion, showing that its function was crucial to infection. Thus, it was possible to endow wild-type and recombinant baculovirus occlusion bodies with ChiA proteins from stably transformed insect cell lines. This opens up potential uses for enhancing viral proteins in IPM with AcMNPV.

## Materials and Methods

### Cells, viruses, and media

*Spodoptera frugiperda* Sf9 cells obtained from American Type Culture Collection (ATCC) were cultured at 27°C as a monolayer in TNM-FH insect medium (SIGMA) with antibiotic-antimycotic solution (GIBCO). *Autographa californica* multiple nucleopolyhedrovirus (AcMNPV) wild type (wt) was obtained from Pharmingen. Wt and recombinant baculoviruses were propagated in Sf9 cells and used to generate transformed insect cell lines.

### Insects

The experiments were conducted using *Rachiplusia nu* larvae and eggs provided by AgIdea (Pergamino) from established insect colonies and maintained at the IMYZA (INTA-Castelar). The insects were reared in trays at 23-25°C in a 70% humidified chamber with a 16:8 light: dark photoperiod, and fed a high-wheat-germ diet until they reached their fifth instar (20 days of age). After the bioassay was performed, the surviving individuals were sacrificed under previous ice-cold anesthesia. Infected individuals’ cadavers were collected for further analysis.

### Recombinant baculovirus obtention

Recombinant baculovirus AcPOLHGFP (polh+) was generated using the Bac-to-Bac methodology according to the supplier’s suggestions (Invitrogen) as previously described (López et al., 2024). The first step was the obtention of the transfer plasmid based on the commercial vector pFastBac-DUAL (Invitrogen). This vector carries a copy of the polh gene in cis (polh+). DH10Bac cells were transformed with the transfer vector, and the recombinant bacmid DNA obtained was confirmed by PCR using M13Fw and M13Rv universal primers and used to transfect Sf9 cells using Cellfectin II® reagent (Invitrogen). Cells were incubated for 5 h at 27°C, after which the transfection medium was replaced with 5 ml of TNM-FH medium. After 5 days, the transfection supernatant was collected. After two passages in cells, the viral stocks were titered by endpoint dilution (O’Reilly et al., 1993) using the Sf9GFP cell line developed in our lab based on the pXXL POLH promoter (Hasse et al., 2015).

### Transfer vectors p_XXL_ChiA and p_XXL_ChiA_NLS_ construction

For the expression of ChiA in stably transformed insect cells, the InsectSelect^TM^ BSD system (Invitrogen) based on the modified commercial plasmid pIB/V5-His-CAT was used, as detailed below. The starting vector for stable transformation of Sf9 lines with the two versions of ChiA was the p_XXL_CAT plasmid, previously obtained from pIBV5HisCAT (López et al., 2010). The sequence of the AcMNPV ChiA enzyme gene was amplified from a bacmid template carrying the complete virus genome in two versions: native sequence without its amino-terminal signal peptide KDEL sequence (ChiA), and another version identical but with the addition of the polyhedrin nuclear localization sequence KRKK (ChiA_NLS_). The oligonucleotides used for amplification, which were designed from the sequence obtained from GenBank under code NC_001623.1, are listed in Table 1.

**Table 1:**
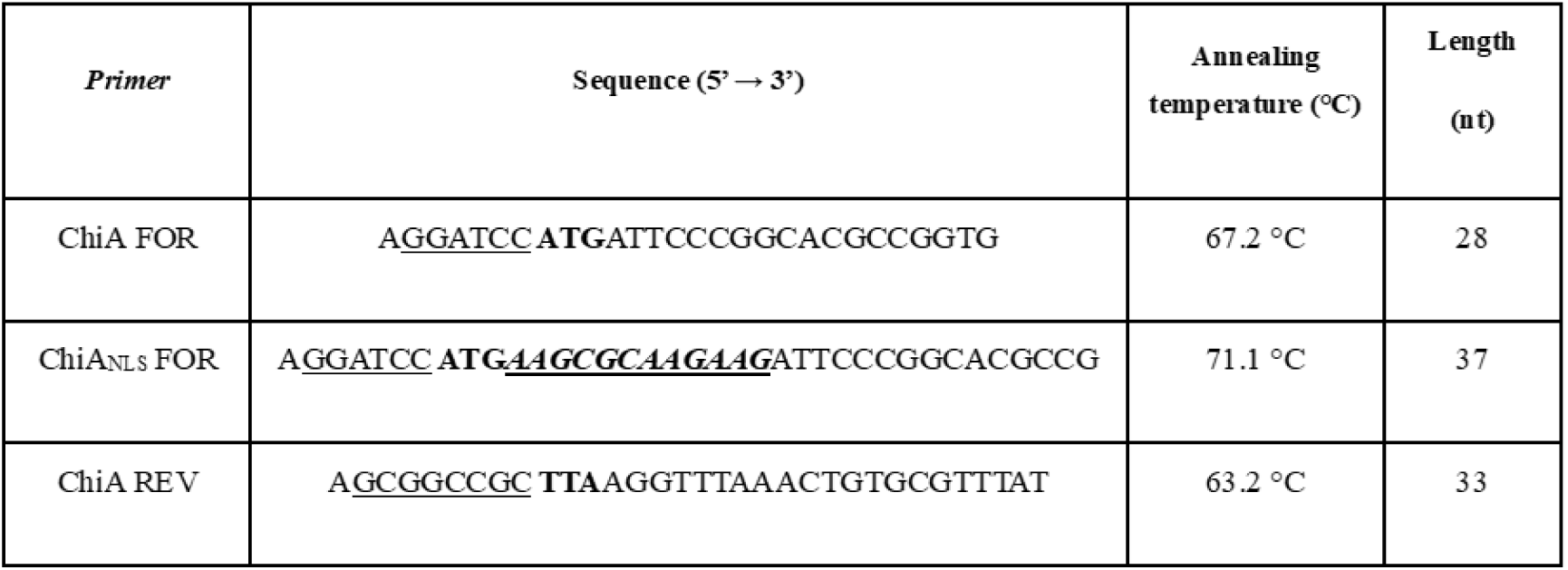
Oligonucleotides used for PCR amplification of the ChiA gene. The cleavage sequences of the restriction enzymes are underlined. The sequence for the translation start codon (AUG) is indicated in bold. The sequence corresponding to the nuclear localization signal (KRKK) is shown in bold italics and underlined. The sequence for the stop codon (UAA) is indicated in bold.

Taq Platinum Pfx enzyme (Invitrogen) was used for the PCR reaction. The sequences amplified and purified in the previous step were reacted with Taq polymerase (GoTaq, Promega) to add 3’ protruding deoxyadenosine ends. and the fragments were then ligated to the commercial vector pCR®2.1-TOPO® (Invitrogen), following the manufacturer’s recommended protocol to obtain vectors TOPO-ChiA and TOPO-ChiA_NLS_. The amplicons ChiA and ChiA_NLS_ were subcloned into p_XXL_ vector using the restriction enzymes *Bam*HI and *Not*I (Thermo Scientific™) to obtain the final transfer vectors p_XXL_ChiA and p_XXL_ChiA_NLS_.

### Stably transformed insect cell lines

For the obtention of the stably transformed insect cell lines, once the identity of the p_XXL_ChiA and p_XXL_ChiA_NLS_ transfer vectors was confirmed, Sf9 cells were stably transfected using Cellfectin II® reagent. Briefly, 2 × 10^6^ cells in 60 mm^2^ culture dishes were transfected with 1 ng of each plasmid using 8 μg of Cellfectin® reagent for each reaction. Experimental controls consisted of: transfected cells without Cellfectin, mock-transfected cells, and untreated cells. In parallel, Sf9 cells were transfected with the pIB/V5-His-GFP plasmid construct to assess the transfection efficiency.

Recombinant cellular clones were selected by blasticidin resistance, starting from a concentration of 60 μg/ml blasticidin previously determined in a cell death curve (López, personal communication). To establish the polyclonal cell lines (Sf9ChiA and Sf9ChiA_NLS_), confluent monolayers were consecutively transferred to larger supports in culture medium and blasticidin at a decreasing concentration with passages, until they were maintained at a constant concentration of 10 μg/ml.

### Cell infection conditions and optical microscopy

Sf9 and Sf9ChiA cells (2 × 10^6^) were seeded per well in six-well chambers for 1 h. Subsequently, infections were performed with the recombinant baculovirus AcPHGFP and AcMNPV wt at a multiplicity of infection (MOI) of five. Four days post-infection (dpi), the cells were visualized under inverted fluorescence microscopy (Olympus LH50A, Olympus, Shinjuku, Tokyo, Japan) with 100X magnification; images of 10 fields were taken for each infection event. The cells were collected in 1X phosphate-buffered saline (PBS; 137 mM NaCl, 2.7 mM KCl, 8 mM Na_2_HPO_4_, and 2 mM KH_2_PO_4_, pH 6.2) after taking the images and processed as samples for Western blot assays.

### Recombinant polyhedra (rOBs) isolation

Sf9 and Sf9ChiA cells (1 × 10^7^) in T175 bottles were infected at an MOI of 5 with the recombinant baculovirus previously obtained. Polyhedra were purified at 5 dpi according to O’Reilly et al. (1993). Briefly, the cells were collected, washed with PBS 1X pH 6.2, and sonicated (three pulses of 30 s at 35% amplitude) in a VCX 500 sonicator (Sonics, OK, USA). Then, the cellular pellet was resuspended in equal volumes of 0.5% sodium dodecyl sulfate (SDS) in glass haemolysis tubes, followed by centrifugation in a low-speed centrifuge (Eppendorf, Hamburg, Germany) at 4000 g for 10 min. The pellets were resuspended in equal volumes of 0.5 M NaCl, followed by centrifugation at 4000 g for 10 min. The pellets containing insoluble material (polyhedra) were finally resuspended in equal volumes of ultrapure water. The isolated polyhedral of each condition were counted in Neubauer chambers and kept at −20°C until use. Statistical analysis of quantification data was obtained by a non-parametric paired t-test using GraphPad Prism version 5.0.0 for Windows, GraphPad software (Boston, MA, USA, www.graphpad.com).

### SDS-PAGE and Western blot assays

The detection of protein incorporation into recombinant polyhedra was performed by sedimenting rOBs, as previously described, dissolved with Na_2_CO_3_ 0.1 M, and resuspended in cracking buffer 1X (120 mM Tris-HCl, pH 6.8, 4% SDS, 0.02 % bromophenol blue, 1.4 M 2-ß-mercaptoethanol, 20% glycerol). The samples were electrophoresed in 12% polyacrylamide gels using Laemmli’s discontinuous buffer system under denaturing conditions and transferred to nitrocellulose membranes for Western blot assays. Detection of ChiA inside polyhedra of different conditions of infection was performed by immunodetection of polyhedra samples isolated from infections with WT, recombinant AcPOLHGFP (PHGFP) baculovirus in two different insect cell lines, Sf9 and Sf9ChiA_NLS_, using a mouse-produced anti-POLH primary antibody and an alkaline phosphatase-conjugated anti-mouse secondary antibody. The Western blot of the same samples was probed with a rabbit-made primary antibody anti-ChiA (1:3000) and an alkaline phosphatase-conjugated anti-rabbit secondary antibody. The expected molecular weight for ChiA is 59 kDa.

### Bioassays

Second instar larvae of *Rachiplusia nu* were starved for 24 h and individually inoculated using the droplet-feeding technique (Hughes and Wood, 1981). Briefly, the larvae were allowed to ingest viral suspensions prepared in 1% sucrose and 0.1% Coomassie Brilliant Blue (phagostimulant solution). Larvae that had consumed the suspension within 5 min were transferred to plastic plates containing fresh virus-free diet and maintained at 25°C. Control larvae were treated similarly, but using water instead of viral suspension. The concentration of OBs for each viral suspension was adjusted based on the mean ingested volume per larva (Kunimi and Fuxa, 1996). *R. nu* larvae were treated with samples of wt and recombinant polyhedra as described in the results section. Three replicate assays of 24 larvae each were performed. Mortality was recorded every 24 h during two weeks. Mortality data were subjected to a one-way analysis of variance (ANOVA) and Tukey’s test (p < 0.01) using GraphPad Prism software.

### Transmission electron microscopy (TEM) of purified OB samples

The purified OBs samples were centrifuged at 12000 rpm for 10 min and fixed in 2% glutaraldehyde in phosphate buffer (pH 7.2–7.4) for 2 h at 4°C. Secondary fixation was performed with 1% osmium tetroxide for 1 h at 4°C. Subsequently, the samples were dehydrated through an ascending ethanol series and embedded in Spurr’s resin. Ultrathin sections (90 nm) were obtained using a Leica EM UC7 ultramicrotome and stained with uranyl acetate and lead citrate. The samples were then examined under a Zeiss EM 109T transmission electron microscope, and images were acquired using a digital Gatan ES1000W camera (LANAIS-MIE. IBCyN, Buenos Aires, Argentina).

## Results

### 1. Stably transformed Sf9ChiA and Sf9ChiA_NLS_ lines allow passive incorporation of ChiA into AcMNPV wt and recombinant OBs

Intending to obtain chimeric polyhedra incorporating the AcMNPV ChiA protein, we addressed the construction of stably transformed Sf9 insect cell lines. In these lines, ChiA expression was induced *in trans* by infection and passively incorporated into the nascent polyhedra. To this end, the AcChiA coding sequence was placed under the regulation of the XXL promoter (Lopez et al., 2010), which encompasses the very strong, infection-inducible polyhedrin promoter, incorporating specific regulatory sequences approximately 2600 bp upstream of the minimal promoter.

To obtain the p_XXL_ChiA vectors, the next step was to amplify the two versions of the proposed ChiA gene. To do this, we first performed an *in silico* study of the gene and amino acid sequence extracted from GenBank. Using Expasy’s SignalP program, it was possible to detect the existence of a signal peptide at the N-terminal end that cleaves at amino acid 17 and, in addition, the KDEL endoplasmic reticulum targeting sequence at the C-terminal end. Based on this characterization, the primers were designed to amplify by PCR two variants of the ChiA gene: one version without signal peptide or KDEL sequence, which was called ChiA, and a second version, the same as the previous one, but with the addition of the nuclear localization sequence present in the polyhedrin gene, KRKK, at the N-terminal end, which we called ChiA_NLS_. This last version was constructed to test whether targeting the synthesis of ChiA to the cell nucleus for passive incorporation into polyhedra is more efficient than when it is synthesized without a localization signal, which is probably destined for the cytoplasm. To add the nuclear localization signal in the ChiA_NLS_ version, the nucleotide sequence corresponding to KRKK extracted from the polh sequence was incorporated into the ChiA amplicon from the primers used in PCR, as described in M&M.

When the stably transformed lines were obtained, a characterization was performed by infecting them with different baculoviruses and studying their evolution and performance in terms of chimeric polyhedra. As mentioned above, according to the sequence chosen for the expression of recombinant AcMNPV ChiA in insect cells, the PM of the protein was estimated to be approximately 60 kDa.

The characterization of the stable lines obtained involved evaluating the effect of infecting these transgenic cell lines with a baculovirus, as a control. This resulted in a band of expression of ChiA in both cell lines, Sf9ChiA and Sf9ChiA_NLS_ (Fig. 1A). As expected, no bands were observed in the cell extracts derived from the stably transformed cell controls (lanes 3 and 4, uninfected), thereby indicating that in the absence of induction, ChiA was not expressed. Furthermore, the employed methodology does not detect endogenous ChiA expression in Sf9 (lanes 1 and 2). Consequently, the expression of inducible ChiA is the only event detectable by this approach. As demonstrated in Figure 1B, for instance, for Sf9ChiA_NLS_, both stably transformed Sf9 cell lines were capable of producing OBs that exhibited a morphology similar to that of the WT. Finally, when polyhedra obtained from WT infections of both Sf9ChiA and Sf9ChiA_NLS_ cell lines were isolated and immunoblotted against ChiA, the highest level of protein incorporation was detected in the Sf9ChiA_NLS_ cell line by chemiluminescence (Fig. 1C). Therefore, the research was continued with that version of the gene.

**Fig. 1:**
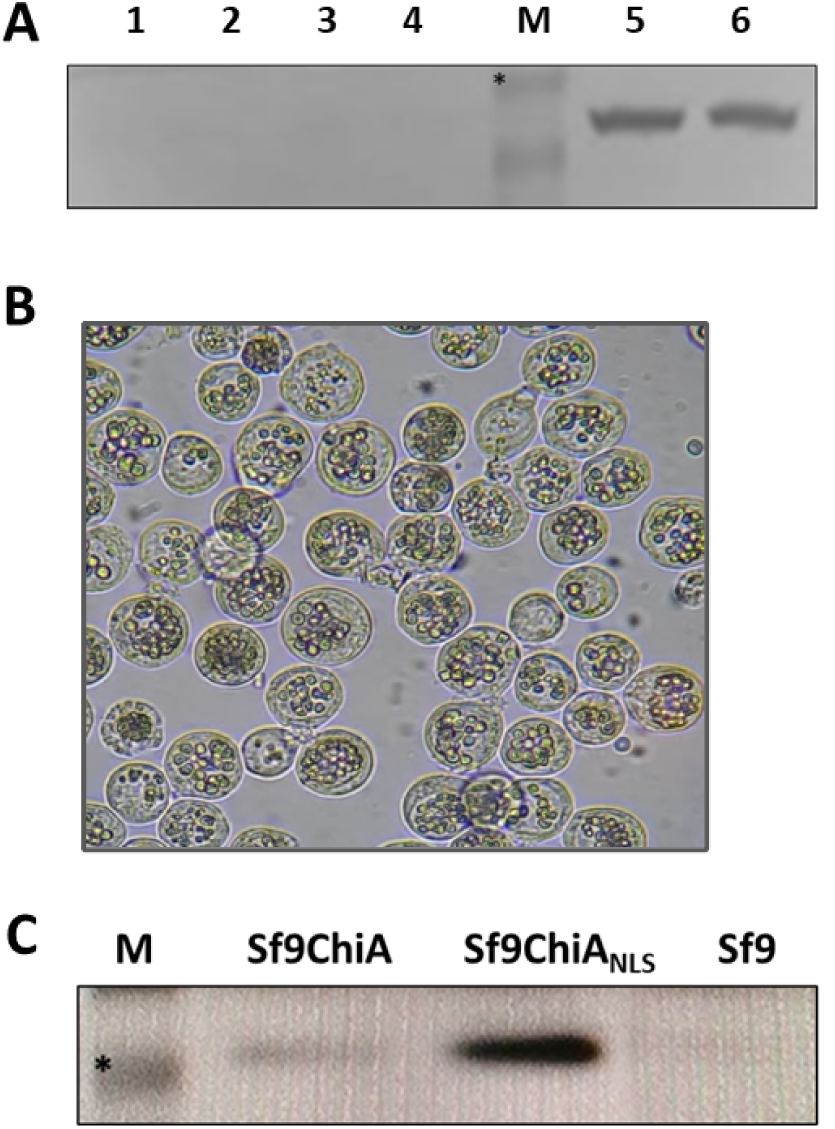
Characterization of stable ChiA cell lines. **A)** Western blot of AcPOLH-baculovirus-infected cell samples performed with a rabbit anti-ChiA primary antibody, revealed with an alkaline phosphatase-conjugated anti-rabbit antibody. 1- Uninfected Sf9 cells 2- Infected Sf9 cells 3- Uninfected Sf9ChiA cells 4- Uninfected Sf9ChiA_NLS_ cells. M: Molecular weight marker. Asterisked band: 70 kDa 5- Infected Sf9ChiA cells 6- Infected Sf9ChiA_NLS_ cells. **B)** Optic microscopy of Sf9ChiA_NLS_ cells infected with WT baculovirus. **C)** Immunodetection of polyhedra samples isolated from infections with WT baculovirus in three different insect cell lines. Using a rabbit-produced anti-ChiA primary antibody and a peroxidase-conjugated anti-rabbit secondary antibody. Revealed by chemiluminescence. M: molecular weight marker. Asterisk: 50 kDa band. 1- WT polyhedra isolated from infection in the Sf9ChiA line 2- WT polyhedra isolated from infection in the Sf9ChiA_NLS_ line 3- WT polyhedra isolated from infection in the Sf9 line.

### 2. Passive incorporation of ChiA

After proving its effectiveness, the cell lines were tested in their ability to passively incorporate ChiA into wild-type and recombinant baculoviruses with polyhedrin fusions. For this purpose, infections were performed in both Sf9 and Sf9ChiA_NLS_ with WT and the recombinant baculovirus AcPOLHGFP (polh+). The baculovirus AcPOLHGFP that incorporates into its genome the reporter gene GFP sequence from *Aequorea victoria* as a fusion to polyhedrin was previously obtained in the laboratory using as a base the commercial plasmid pFastBac-DUALl, which presents two promoters: pPOLH and pP10, and has, in addition to the fusion protein, an additional copy of wild-type polyhedrin to ensure the correct formation of occlusion bodies. The transfer vector construct used to obtain the recombinant baculoviruses is shown in Fig. 2A.

**Fig. 2.**
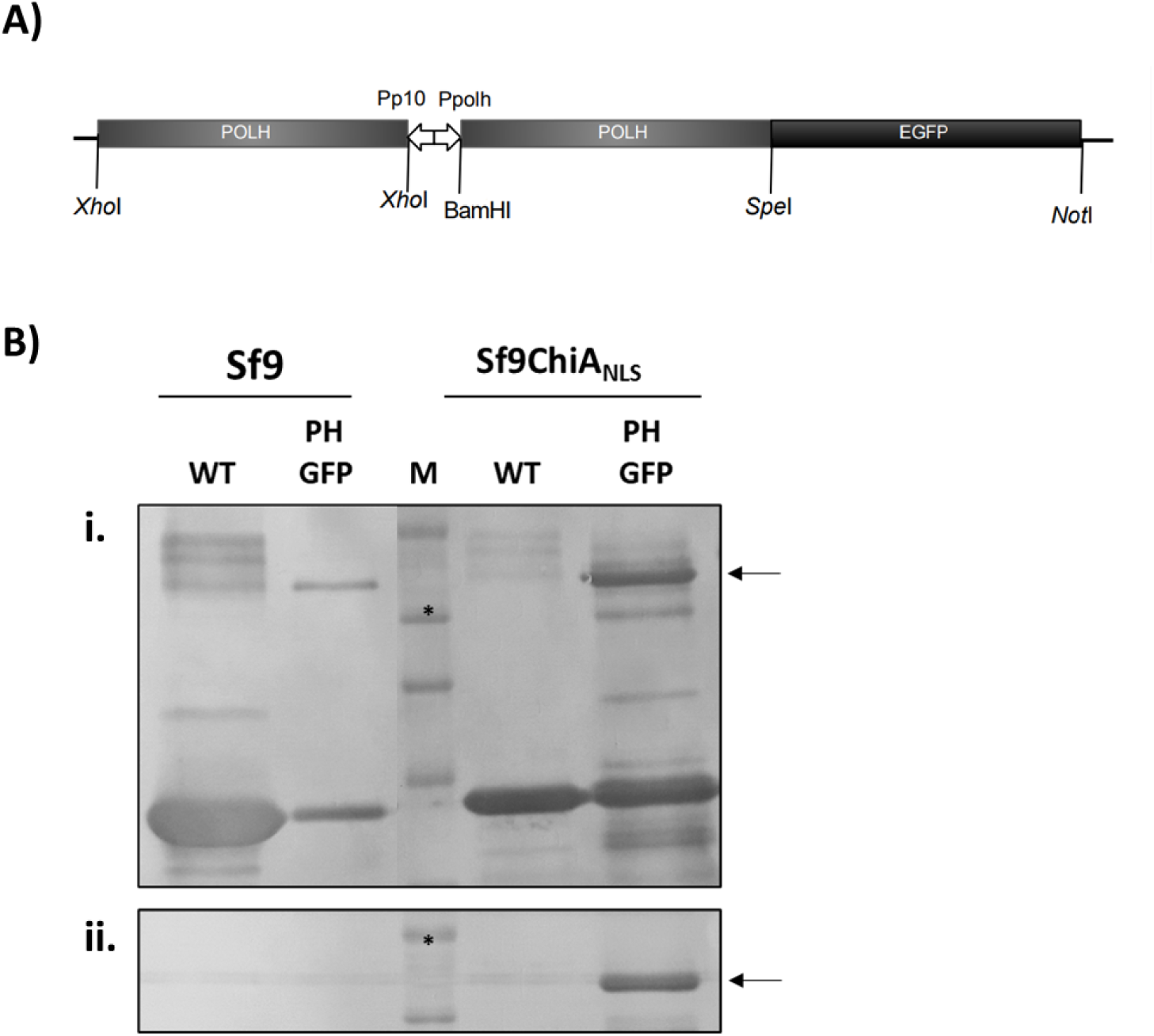
Detection of ChiA within polyhedra under different infection conditions. **A)** Schematic graph of the cassette cloned into the transfer vector used in the construction of recombinant baculovirus AcPOLHGFP, based on commercially available pFastBac-DUAL (Invitrogen). **B) i.** Immunodetection of polyhedra samples isolated from infections with WT and recombinant AcPOLHGFP (PHGFP) baculovirus in two insect cell lines, Sf9 and Sf9ChiA_NLS_. Using a mouse-produced anti-POLH primary antibody and an alkaline phosphatase-conjugated anti-mouse secondary antibody. M: Blue Plus (TRANSGene Biotech) molecular weight marker. The asterisk marks a band of 50 kDa. **ii.** Western blot of the same samples as in i, but using a rabbit anti-ChiA primary antibody and an alkaline phosphatase-conjugated anti-rabbit secondary antibody. Asterisked band: 70 kDa. Arrows show the incorporated recombinant proteins.

Polyhedra from two infection conditions were treated with 0.5% SDS and washed with sequential centrifugation steps and then dissolved with Na_2_CO_3_, before the proteins were resolved on an SDS-PAGE. These sequential washes minimize external protein contamination, and carbonate dissolution selectively solubilizes the polyhedrin matrix to enrich intramatrix components. The gel was transferred to a nitrocellulose membrane, and first, for the detection of the proteins fused to POLH, the membrane was probed with a mouse-produced anti-POLH primary antibody and an anti-mouse secondary antibody conjugated to alkaline phosphatase (Fig. 2Bi). In parallel, the same samples were run and transferred in the same manner, and ChiA protein in each sample was detected with the anti-ChiA antibody used in previous assays and a secondary antibody conjugated to the enzyme alkaline phosphatase to be revealed by colorimetric substrates (Fig 2Bii).

With this assay, we confirmed the obtention of chimeric polyhedra containing two recombinant proteins in part of their matrix: GFP and ChiA. As expected, the fluorescent protein was actively incorporated through fusion with POLH (see Fig. 2Bi, arrow). Note that ChiA was detected in the same lane (Fig. 2Bii, arrow), confirming that it was passively endowed within the polyhedra of the recombinant baculovirus. In contrast, ChiA could not be detected in the wild-type baculovirus polyhedra using this technique (colourimetric alkaline phosphatase substrates).

### 3. AcChiA improved the levels of infectivity of the wt and recombinant baculovirus

To assess whether the incorporation of ChiA into polyhedra increased baculovirus infectivity, time-dependent lethality bioassays were conducted on larvae of the lepidopteran species *Rachiplusia nu*. Stable lines were infected with different baculoviruses to obtain viral inocula, and the assays were performed as described in the Materials and Methods section.

Figure 3 shows the results, indicating significant differences only between the treatments with and without ChiA incorporated in the polyhedra for AcMNPV WT and AcPOLHGFP baculoviruses.

**Fig. 3.**
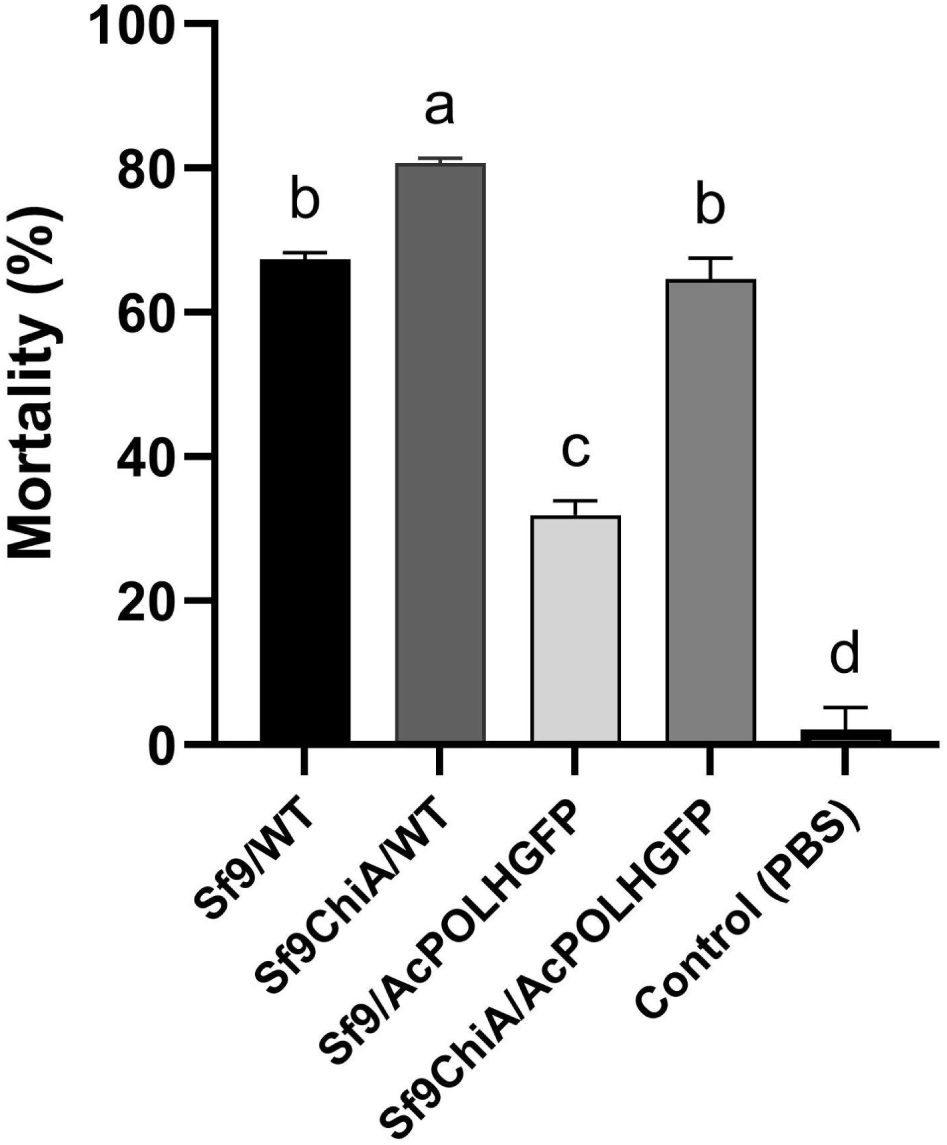
Mortality percentages recorded on *R. nu* larvae infected with recombinant polyhedra. WT: AcMNPV wild type. Control: without inocula. Different letters indicate significant differences between mortality rates (p > 0.05).

The results of this study indicate that the incorporation of ChiA into the polyhedra led to an augmentation in the natural infectivity of the viruses under investigation. The analysis of variance (ANOVA) revealed an increase of 13.35% between these conditions (Fig. 3 Sf9/WT vs Sf9ChiA/WT). However, it was found that recombinant polyhedra exhibited a higher propensity to incorporate ChiA protein into their structure in comparison to the WT sample, showing an increase of 32.75% in mortality (Fig 3. Sf9/AcPOLHGFP vs Sf9ChiA/AcPOLHGFP). The presence of this protein in the chimeric polyhedra may contribute to the observed increase in infectivity. In the case of wild-type (WT) viruses, incorporation was not detected; the increase in infectivity is less evident than that observed for the recombinant AcPOLHGFP.

### 4. Recombinant baculoviruses exhibited a significant reduction in ODV occlusion

To ascertain the factors that could be affecting the mortality rates observed in bioassays, and also to confirm that the incorporation of ChiA into the polyhedra structure did not alter the normal morphogenesis of OBs, an investigation was conducted into the composition of WT and recombinant OBs, purified from an Sf9ChiA cell line infected with both baculovirus AcPOLHGFP and WT. The present investigation entailed the observation of the presence of ODVs in purified polyhedra ultrathin sections by TEM. The findings confirmed the hypothesis that the incorporation of ChiA did not alter the morphology of the polyhedra in any of the samples and also that the incorporation of ODV into polyhedra was notably reduced for AcPOLH-GFP compared to WT (see Figure 4). This low incorporation of ODV into polyhedra may explain the low mortality rate of AcPOLH-GFP observed in bioassays (see Figure 3). The images selected were illustrative of a broader set of images that shared analogous characteristics.

**Fig. 4.**
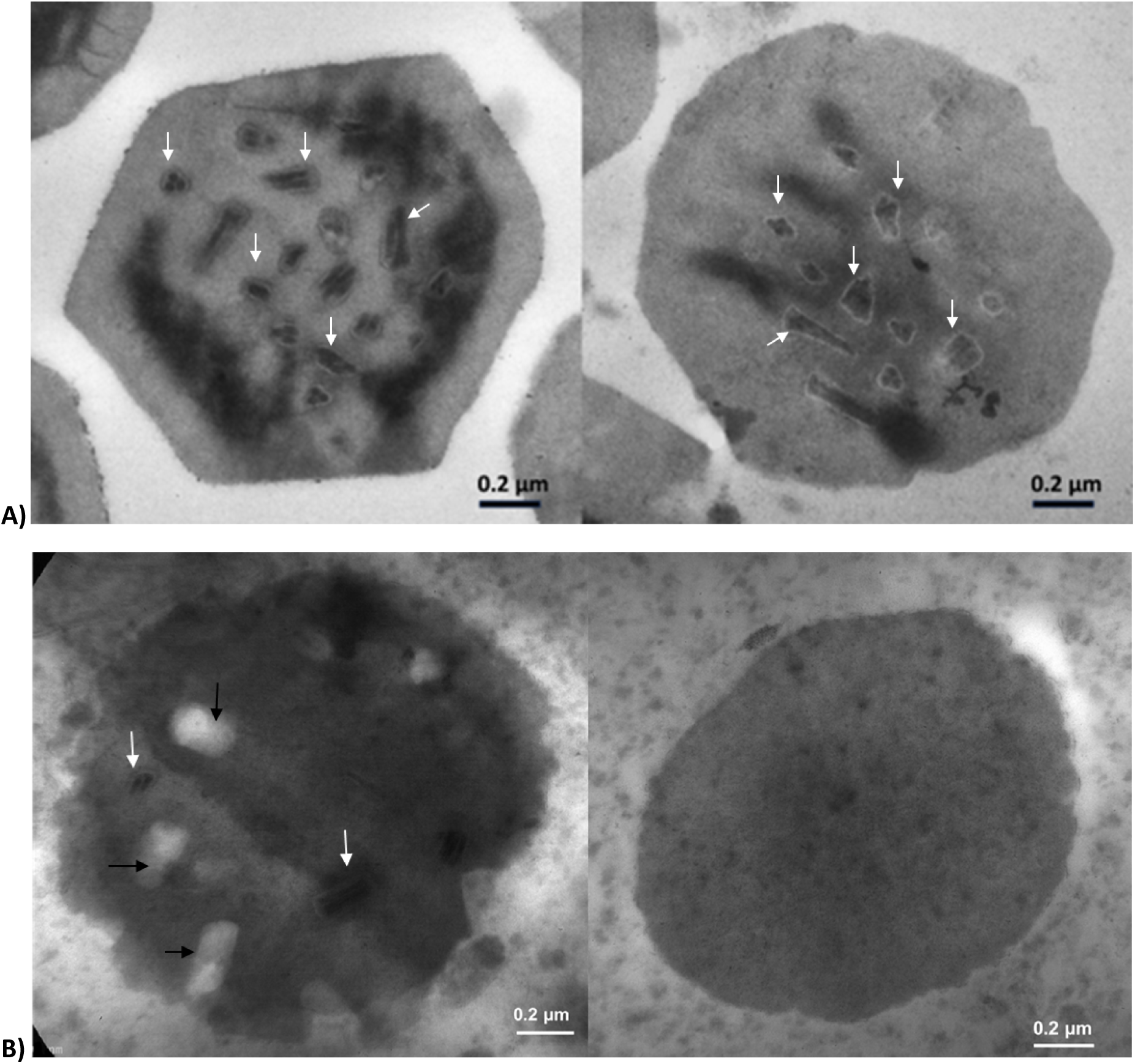
Transmission electron microscopy (TEM) images of microtome-ultrathin sections of purified Sf9ChiA/AcMNPV WT (A) and Sf9ChiA/AcPOLHGFP (B) polyhedra samples. The white arrows show ODV’s presence inside the polyhedra matrix. The black arrows indicate the probable gaps left by ODVs that are not observed in the samples of polyhedron sections obtained from infecting the lines with recombinant baculoviruses.

TEM observations demonstrate that WT baculovirus exhibited normal occlusion in the Sf9ChiA cell line (indicated by white arrows, Fig. 4). However, few ODVs were observed to occupy the protein matrix of the recombinant polyhedra in the Sf9ChiA cell line. This was evident in thin sections of polyhedra, where no ODVs were visible, or where spaces that could have been occupied by ODVs were evidenced as white gaps (black arrows, Fig. 4B). This fact could explain the reduction in infectivity provoked by the recombinant baculovirus, which entails the introduction of a fusion protein into the POLH sequence, resulting in the destabilisation of the crystalline structure of the polyhedron, and that its recomposition in the case of inocula obtained by infection of Sf9ChiA (see Figure 3), could be attributed to the incorporation of ChiA into the polyhedra matrix.

### Concluding remarks

This study demonstrated that the incorporation of ChiA into the polyhedra matrix could be achieved through a passive process involving stably transformed insect cell lines that contribute the protein *in trans*. These cell lines were capable of embedding both WT and recombinant polyhedra with that viral protein, resulting in a substantial augmentation in their mortality rates. In the case of recombinant polyhedra, an additional observation was made: namely, a notable reduction in ODV. In this instance, it was found that ChiA was acting as an infectivity potentiator. All these findings are important for the design of new control strategies in IPM using AcMNPV.

## Discussion

Integrated pest management programmes are currently the most widely accepted strategy for reducing the use of chemical insecticides. Biopesticides play a crucial role in this framework, complementing and, in some cases, replacing conventional management methods. Baculoviruses are considered an effective and safe option within this group, but require improvement to reduce infectivity times and to compete with chemical pesticides. Baculovirus-based insecticides are attractive owing to several advantages over other products. Indeed, they are environmentally innocuous and harmless to humans and animals, and they have long-lasting control effects on the target insects (Haase et al., 2015). Currently, the engineering of baculoviruses for use as bioinsecticides is focused on expressing insect-specific or viral genes that modify the physiology of the target insect (Kroemer et al., 2015). In addition to using recombinant baculoviruses that express soluble proteins to enhance their action, another possible strategy to improve their insecticidal capacity is the generation of baculoviruses that incorporate recombinant proteins into their occlusion bodies. Jung and colleagues were among the first to attempt the use of recombinant OBs (rOBs) as bioinsecticides. They developed polyhedra that incorporated the *Bacillus thuringiensis* (Bt) Cry1Ac toxin and GFP as a marker. Nevertheless, in the last decade, most of the attention has been turned towards the use of OBs as a promising biotechnological tool (Chang et al., 2003; Jung et al., 2012). Given the wide range of genetic modifications tested on baculoviruses, the most effective approach to enhancing their killing efficacy compared to wild-type baculoviruses seems to be the addition of a toxin to occlusion bodies (Kamita et al., 2010).

This work aimed to incorporate viral proteins into AcMNPV occlusion bodies that can disrupt the peritrophic membrane of insects. This has the potential to improve the infectivity of baculoviruses. AcChiA, a viral enzyme responsible for the hydrolysis of chitin produced by many organisms, including crustaceans, fungi, and insects, could be a potential insect control protein, as its substrate, chitin, is a component of the insect peritrophic membrane (PM) (Kramer and Muthukrishnan, 1997). ChiA of AcMNPV could be expressed in *E. coli* bacteria and administered to *Bombyx mori* as a bioinsecticide, reaching levels of mortality at 60 μg ChiA/g larva (Rao et al., 2004). In earlier research, it was demonstrated that a variety of strategies can be employed to incorporate proteins into OBs using fusions to POLH, fragments of POLH, and the contribution of transformed insect cell lines (López et al., 2024). In the context of biological control, it is imperative to prioritize the utilization of wild viruses over recombinant viruses. Consequently, the strategy that emerges as the most promising is that of *in trans* incorporation of proteins into OBs, which significantly minimizes the reliance on recombinant viruses in field applications.

To incorporate proteins into OBs through stable insect cell lines, a nuclear localization signal (NLS) is required. The chosen signal was polyhedrin’s KRKK. The aim of constructing the Sf9ChiA_NLS_ line was to analyse the effect of the incorporation of the nuclear localization signal POLH on the amount of ChiA in the nucleus and to test whether it influences the levels of its passive incorporation into polyhedra. The possibility of passive incorporation of proteins into polyhedra was tested in BmMNPV by overexpressing a reporter protein with the same viral vector, which was subsequently found to be part of the composition of OBs (Xiang et al., 2012). Subsequent studies in the same species reinforce the flexible and plastic properties of the polyhedron matrix to incorporate or embed other proteins into its structure (Guo et al., 2018). Against this background, it was expected that overexpression of ChiA by stable cell lines as a new strategy for overexpression would lead to its passive incorporation into polyhedra.

The ChiA-transformed cell lines were obtained based on a plasmid previously developed in our group carrying a polyhedrin promoter optimised by the addition of enhancer sequences (pXXL) used in the construction of other stably transformed lines, which have been shown to produce high levels of expression and to be inducible by baculoviral infection (López et al., 2010; Plastine et al., 2024). These characteristics, together with the fact that it is a very late promoter, which would allow the expression of ChiA to coincide with the morphogenesis of the polyhedra, explain the choice of this system for the strategy proposed in this work. Both cell lines, Sf9ChiA without a nuclear localization signal and Sf9ChiA_NLS_ with the signal in the N-terminal end, were obtained and characterized. The transgene was expressed only when the Sf9ChiA and Sf9ChiA_NLS_ cell lines were infected with baculovirus. Infections were performed at matched MOI, and OBs were purified identically across producer lines; moreover, OB morphology remained similar (Figure 1B), mitigating concerns about gross structural differences. Additionally, the ChiA enzyme expressed by the stable lines obtained could be passively incorporated into the OBs generated when infected with a wild-type baculovirus (see Figure 1C). With this experiment, we could notice that the detection of ChiA was method-dependent, as a stronger signal was recorded for recombinant infections, while detection in WT was weaker and sensitive to the chemistry used. The result is consistent with previous observations regarding the structure of POLH and its ability to accommodate proteins other than baculoviral proteins within polyhedra (Chiu et al., 2012). This study did not directly visualize ChiA within the occlusion body matrix; immunogold TEM or protease-protection assays will be required to establish definitive localization. In the case of this recombinant baculovirus, no morphological changes in OBs were observed, but a notable reduction in ODV occlusion was detected, probably due to recombinant protein fusions to POLH. As previously stated, further investigation into this matter would be worthwhile to establish the reasons for the decrease in ODV in this system.

Despite the lower level of ChiA detection intensity observed in the Sf9ChiA cell line, it can be concluded that this protein was successfully incorporated into AcMNPV polyhedra even without a core-targeting signal. The observed difference in incorporation levels between the two lines may suggest a positive effect of adding the KRKK nuclear localization sequence on the amount of ChiA present in wild-type polyhedra. However, further studies are required to confirm this. Comparing different strategies, the incorporation of more ChiA protein was enabled by the recombinant structure of chimeric polyhedra conformed by trimers of POLHGFP mixed with trimers of wt POLH. This finding was confirmed by WB assays of dissolved OBs (Fig. 2B). This finding is consistent with previous reports demonstrating the flexibility of recombinant polyhedra structures to facilitate the incorporation of heterologous proteins (Sampieri et al., 2015; Guo et al., 2018; Diez et al., 2018). Subsequent research determined that recombinant baculoviruses of this nature, which actively incorporate proteins into OBs by fusing to POLH (López et al., 2018), exhibit ODV occlusion deficiencies. This phenomenon can be added to the naturally occurring amino acid substitutions in polyhedrin, which have also been shown to produce a variety of phenotypic changes to OBs. A significant proportion of these substitutions are classified as single-point mutations, which occur within specific structural domains of the protein. The phenotypic changes range from large, cuboid polyhedra, which occlude no or few virions (Lin et al., 2000; Ribeiro et al., 2009; López et al., 2011), to overall changes in polyhedra shape (Cheng et al., 1998; Ji et al., 2010). In the present study, the utilization of TEM was employed to confirm that recombinant BVs incorporating fusions to POLH into OBs exhibited reduced ODVs in comparison to WT BVs infecting the same cell lines. In addition, the aforementioned deficiencies were preliminarily quantified using quantitative PCR (qPCR), thereby corroborating the results (data not shown). These deficiencies may result in a reduction of their fitness compared to WT, thus necessitating further study, which should involve a comparison of these baculoviruses with others. This would assist in determining whether such deficiencies are attributable to polyhedrin structure failures, misfolding, or other issues about OB morphogenesis. It is reasonable to hypothesize that studying this phenomenon would be facilitated by utilizing diffraction equipment specifically designed for analyzing nanoparticles.

In the bioassays, OBs produced in ChiA-expressing lines were associated with higher larval mortality than OBs produced in Sf9, consistent with a ChiA-mediated effect; it was observed that the incorporation of ChiA into wt polyhedra increased the mortality of WT baculoviruses by 13.35%, while enhanced mortality of chimeric polyhedra was observed in 32.75%. As illustrated in Figure 3, the incorporation of recombinant AcPOLHGFP into the polyhedra resulted in a decline in mortality rates. However, when considering the recombinant baculovirus AcPOLHGFP, it becomes evident that ChiA’s contribution was determinant in potentiating the infectivity of these viruses. Given prior demonstrations that AcChiA disrupts the peritrophic membrane (Rao et al., 2004), the increased mortality observed here is consistent with enhanced primary infection via PM poration. Direct measurements of peritrophic membrane permeability and ultrastructure were not performed in this study; future work might be performed with fluorescent tracer assays and midgut TEM to confirm the mechanism of action.

In conclusion, this work presents evidence of high levels of non-directed incorporation of a protein with proven baculovirus oral infectivity-enhancing action into AcMNPV OBs, and has not been reported before. Although the potential of this development has yet to be analysed in the field, it offers a promising advantage in terms of application using wild-type baculoviruses. This would avoid the difficulties associated with the use and release of genetically modified baculoviruses as bioinsecticides. Furthermore, the strategy facilitates the incorporation of additional recombinant proteins through the utilization of recombinant baculoviruses. This approach can be employed for other applications of chimeric OBs (López et al., 2018). Consequently, passive incorporation of ChiA into AcMNPV OBs using an *in trans* strategy emerges as a potential application for IPM using this baculovirus.

